# An automated microfluidic platform integrating functional vascularized organoids-on-chip

**DOI:** 10.1101/2021.12.29.474327

**Authors:** Clément Quintard, Gustav Jonsson, Camille Laporte, Caroline Bissardon, Amandine Pitaval, Nicolas Werschler, Alexandra Leopoldi, Astrid Hagelkrüys, Pierre Blandin, Jean-Luc Achard, Fabrice Navarro, Yves Fouillet, Josef M. Penninger, Xavier Gidrol

## Abstract

The development of vascular networks on-chip is crucial for the long-term culture of three-dimensional cell aggregates such as organoids, spheroids, tumoroids, and tissue explants. Despite the rapid advancement of microvascular network systems and organoid technology, vascularizing organoids-on-chips remains a challenge in tissue engineering. Moreover, most existing microfluidic devices poorly reflect the complexity of *in vivo* flows and require complex technical settings to operate. Considering these constraints, we developed an innovative platform to establish and monitor the formation of endothelial networks around model spheroids of mesenchymal and endothelial cells as well as blood vessel organoids generated from pluripotent stem cells, cultured for up to 15 days on-chip. Importantly, these networks were functional, demonstrating intravascular perfusion within the spheroids or vascular organoids connected to neighbouring endothelial beds. This microphysiological system thus represents a viable organ-on-chip model to vascularize biological tissues and should allow to establish perfusion into organoids using advanced microfluidics.

## Introduction

The ability to vascularize organoids remains a challenge in the field of tissue engineering. Indeed, most tissues exceeding 400 *μ*m in thickness need a functional vasculature to ensure a sufficient supply of nutrients and oxygen, as well as the ability to remove carbon dioxide and cellular waste products, preventing the formation of necrotic inner cores^1^. While several approaches have been used to engineer microvascular networks on-chip, the self-organization of endothelial cells is a preferred way because it closely mimics *in vivo* vasculogenesis and angiogenesis processes^2^. Self-organization can be achieved by seeding endothelial cells and supportive cells in a hydrogel, providing structural matrix and biochemical support to the embedded cells^3^. Much progress has been made in generating perfusable vascular networks on a chip, either as a tissue on its own^4–6^, or by incorporating spheroids such as micro-tumours or pancreatic islets^7–11^. Compared to spheroids, organoids offer the advantage of recapitulating complex 3D organ-specific structure and function. However, they are usually cultured under static conditions where they lack vasculature. Recently, significant efforts have been devoted to the development of vascularized complex organoids *in vitro*^12,13^. However, to the best of our knowledge, no *in vitro* system has shown intravascular perfusion within organoids through functional anastomosis with a neighbouring endothelial network.

Current microfluidic chip designs primarily consist of a central microchamber, bound by two lateral perfusion microchannels. The cell-containing gel is pipetted into the central microchamber, and micropatterned ridges or posts have to be engineered to prevent leakage in the perfusion channels^14–16^. Capillary burst valves have been designed, based on geometric logic, to prevent such leakage^17,18^. However, since fluid handling in such designs is delicate, special and technically challenging loading techniques are often required to slow down the polymerisation of the hydrogels^19–21^. Most chip designs also do not provide an automated trapping mechanism of the biological object of interest. For instance, Nashimoto et al.^20^ manually placed a model spheroid into a specific well, micromachined in a chip. Phan et al.^22^ placed micro-organs and micro-tumours in a microvascular network chip but failed to precisely control their location.

Organ-on-chip technologies have emerged over the last decade with great promise. They are powerful tools to assess basic mechanisms of physiology and disease, improved toxicity testing and screening of new drugs^23^. However, it has become clear that significant efforts are needed to provide robust and accessible ways to generate vascularized organs- and organoids-on-chip^24,25^. Here, we report an innovative organ-on-chip device based on a serpentine microchannel architecture that allows for automated and precise location of spheroids or organoids. The reliability of our system was validated using spheroids generated from fibroblasts and endothelial cells as well as 3D human blood vessel organoids generated from human-induced pluripotent stem cells (iPSCs)^26^. Importantly, we demonstrate effective anastomosis and controlled perfusion of the vascular organoids providing a method for organ-on-chip vascularization that can be adapted to a multitude of other 3D cell aggregates; such as organoids, tumoroids, spheroids, or tissue explants.

## Results

### Design of a serpentine-shaped microfluidic device for precise entrapment of organoids

We set out to design a microfluidic device that is user-friendly, robust and automated. A microfluidic chip was fabricated using Cyclic Olefin Copolymer (COC), a material that offers long-term robustness, is suitable for mass production, has desired optical qualities for imaging, and low attachment of chemicals^27^. The microchannel architecture allowed us to precisely and reproducibly trap an organoid to a predefined location consisting of a U-cup shaped restriction (Fig 1. a,b). The chip was designed based on existing hydrodynamic trapping principles^28^, but with some modifications in consideration of the organoids’ diameters and loading processes (Supplementary Figs. 1,2 and Supplementary Note 1).

**Fig. 1.**
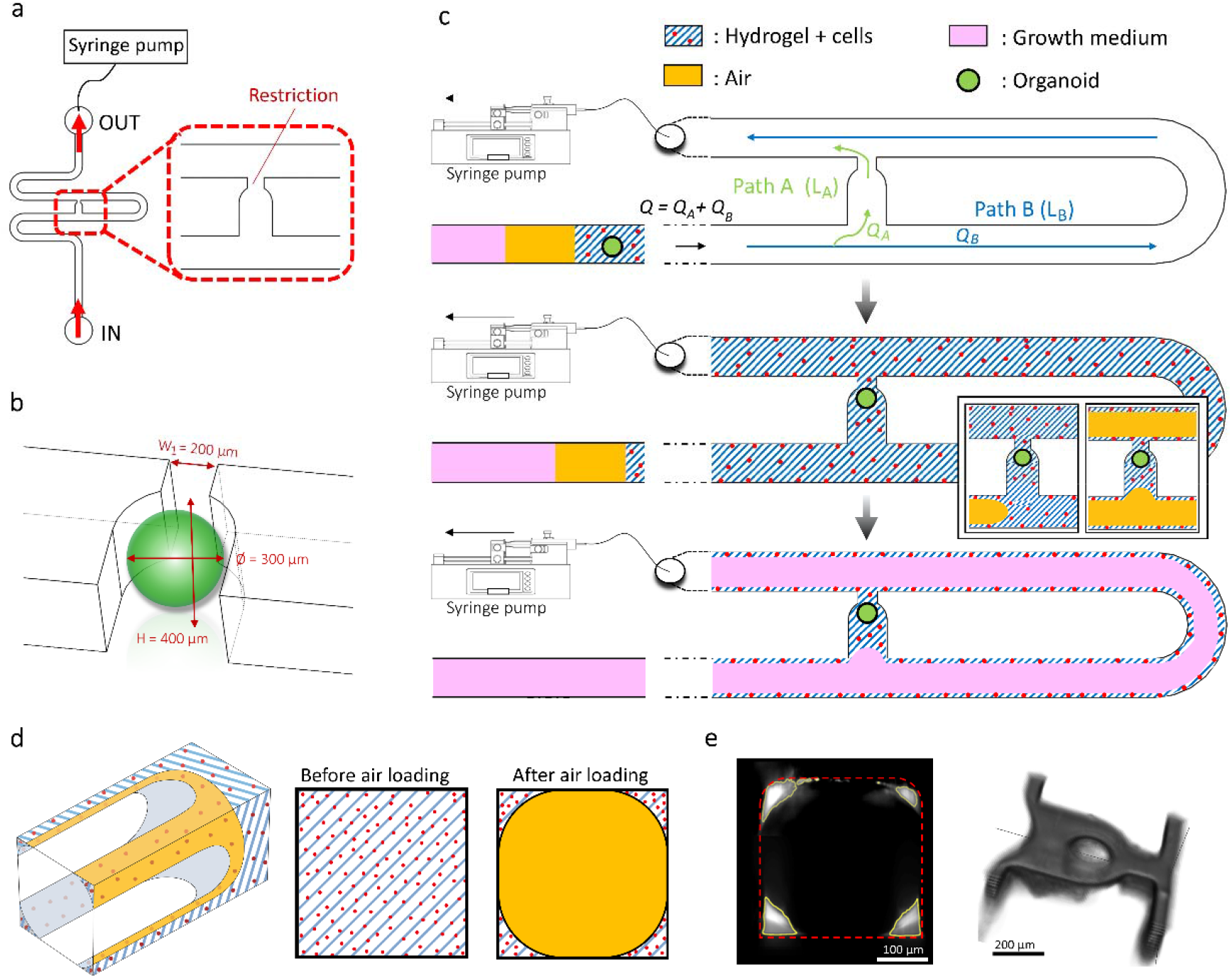
Device design and overview of organoid and cell configurations. **a,** Top view of the microfluidic device. A syringe pump was connected to the outlet of the channel to introduce fluids perfusion. **b,** Schematic three-dimensional view of the U-cup shaped area functioning as a trap. Here, the trap site is exemplary occupied by a cell aggregate of 300 μm in diameter. **c,** Schematic diagram showing an overview of the loading process. Initially, the hydrogel containing an organoid and HUVEC cells was introduced. Before polymerization of the hydrogel, air was introduced to position the hydrogel and the HUVEC cells. Finally, growth medium was introduced for continuous perfusion of the microfluidic chamber and trapped organoid. **d,** Schematic 3D and cross-sectional views of the microchannel showing the air loading process and associated hydrogel deposition. **e,** Experimental cross-sectional view (left) of the microfluidic channel showing the hydrogel deposition in the trap and in the channel’s corners and 3D rendering (right), taken with an in-house light sheet fluorescence microscopy set-up.

Microfluidic trapping through differential hydrodynamic resistance has been proven to be highly efficient^29,30^ and generates low shear stress in the U-cup trap once it is occupied^31,32^. The organoid, embedded in a fibrin hydrogel, was precisely positioned in the trap site without any apparent morphological alteration (Supplementary Fig. 3). After the trapping of the organoid and surrounding endothelial cells, air was injected to push the hydrogel toward the exit of the microfluidic channel. After hydrogel polymerization, the continuous microfluidic perfusion with growth medium was established (Fig. 1c). Both the hydrogel and the air were injected at *Q* = 300 μl/min. Of note, the hydrogel remained inside the trap site due to capillarity, effectively surrounding the organoid, and minimizing organoid contact with the microchannel walls resulting in a permanent organoid encapsulation inside the trap site^33^. Because of the Landau-Levich-Bretherton effect^34^, a thin layer of hydrogel also remained along the square profile cross-section of the microchannels after injection of air (Fig. 1d). We used this property as a way to provide endothelialization of the serpentine channel with human umbilical vein endothelial cells (HUVEC). Experiments using an adapted in-house light sheet fluorescence microscopy set-up were conducted to observe the three-dimensional structure of the hydrogel deposition near the trap site and along the main serpentine microchannel (Fig. 1e and Supplementary Fig. 4). Thus, we have designed a serpentine geometry chip for precise and controlled entrapment of organoids in endothelialized channels.

### Establishment of interconnected endothelial networks

To interrogate the usefulness and biological relevance of our design, we set out to develop endothelial networks in the newly designed microfluidic chips. We used cell aggregates consisting of fibroblasts and GFP (green) labelled HUVEC cells, termed here mesenchymal spheroids, which were seeded into the microfluidic channels. We first examined the effects of fluid flow on the differentiation of these spheroids into vessel-like structures by culturing them alone under static (with media change every day) or flow conditions (Fig. 2a). We observed an enhanced formation of endothelial networks under dynamic perfusion, with a significant 4.4-, 5.0-, 4.8- and 6.5-fold increase in the number of junctions, number of segments, total segment length and number of meshes respectively, as compared to static conditions (Fig. 2b).

**Fig. 2.**
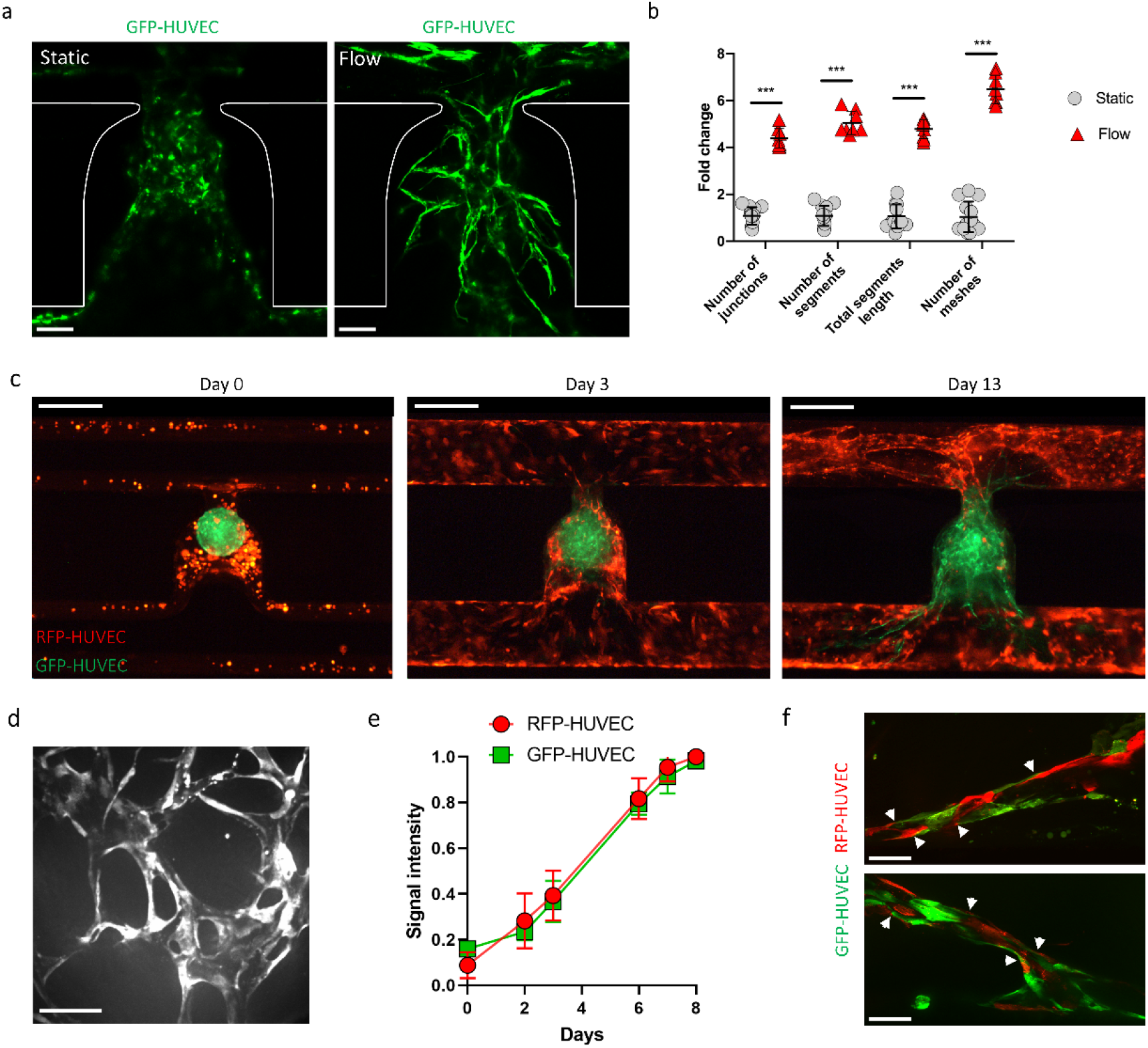
Angiogenic sprouts of RFP-labelled HUVEC (red) cells in the fibrin hydrogel from the main channel anastomose with GFP-labelled HUVEC (green) cells present in the mesenchymal spheroid. **a,** Confocal z-stack maximum intensity projection renderings for GFP-HUVEC cells of the mesenchymal spheroids cultured alone under static or flow conditions. Representative images are shown at day 8 after seeding. **b,** Angiogenesis Analyzer outputs of the abundance and vessel lengths of the endothelial networks, reported as a fold change relative to the static condition. Data points indicate results from n = 13 and 8 (static and flow conditions, respectively) independent microchannels from n = 3 different experiments. **c**, Time resolved evolution of the endothelial network. Gel embedded RFP-HUVEC cells in the main channel are shown in red, GFP-HUVEC cells from the mesenchymal spheroids are shown in green. Note the formation of structured endothelial networks over time that appear to be stable until the end of the observation period (day 13 after seeding). **d**, Confocal z-stack maximum intensity projection of the three-dimensional endothelial network. Image is representative of experiments from n = 12 independent microchannels. **e**, Quantification of RFP-HUVEC and GFP-HUVEC cell proliferation. Intensity signals were normalized by the maximum intensity signal of each image series, corresponding to the RFP-HUVEC and GFP-HUVEC cells covered areas of analyzed microfluidic channel over time, as shown in (**c**) (n = 6 independent microchannels). **f**, Anastomosis between RFP-HUVEC and GFP-HUVEC cells indicated by white arrows. Scale bars, 100 μm (**a**), 400 μm (**c**), and 50 μm (**e**,**f**). Data represents mean ± s.d. Statistical significance was attributed to values of *P* < 0.05 as determined by unpaired t test (two-tailed). \*\*\**P* < 0.001. No significant differences were found in cell proliferations as determined by a two-way ANOVA with Šidák multiple comparisons tests.

Since the hydrogel in the microchannels contains HUVEC cells, we sought to establish connections between the HUVEC seeded serpentine microchannel and the trapped mesenchymal spheroids. The HUVEC endothelial cells of the spheroids expressed GFP (green) and the HUVEC endothelial cells suspended inside the gel prior to injection expressed RFP (red), allowing for direct visualization of these distinct cell populations and to explore interactions between them. On day 0, the mesenchymal spheroids were introduced into the chip and trapped in the correct locations, where they maintained their spherical shape. On day 3, we observed an initial organization of the endothelial cells (Fig. 2c), and by day 7 a three-dimensional endothelial network was formed (Fig. 2d). We also determined endothelial cell proliferation over time in the microfluidic channels near the trap site by quantifying the area of the image covered by the HUVEC cells (Fig. 2e). Due to the presence of fibroblasts in the mesenchymal cell aggregates, the GFP-labelled HUVEC (green) cells were found to spread rapidly. RFP-labelled HUVEC (red) cells initially located in the corners of the main channel, also proliferated and coated the walls of the microchannels.

Next, we examined whether the angiogenic sprouts from the cells suspended in the gel and endothelium formed in the mesenchymal spheroids connected to form a functional network. Intriguingly, we observed spontaneous anastomosis between the RFP-HUVEC endothelial bed and the endothelium from the mesenchymal spheroids, establishing an interconnected network (Fig. 2f). Thus, the serpentine geometry chips we designed allowed us to establish anastomosed endothelial networks.

### Functionality of the endothelial network

To demonstrate the functionality of the interconnected endothelial network, we performed perfusion assays using red fluorescent microbeads. To visualize possible flow through all possible paths within the network, beads with a 1 μm diameter were injected at a flow rate of *Q*_+_ = 10 μl/min to the organ-on-chip on the day 13 of culture (Fig. 3a, Supplementary Video 1). Importantly, the majority of the beads travelled through the endothelial network along the main perfusion direction (path A). As expected, some beads also perfused through the secondary pathway (path B). Of note, as the experiment was done at a constant flow rate, the velocity of a microbead was inversely proportional to the cross-sectional area it flowed through, hence the velocity decreased in regions without endothelium. This resulted in an intense fluorescent signal on the bottom left and top right corners of the imaged area (Fig. 3a).

**Fig. 3.**
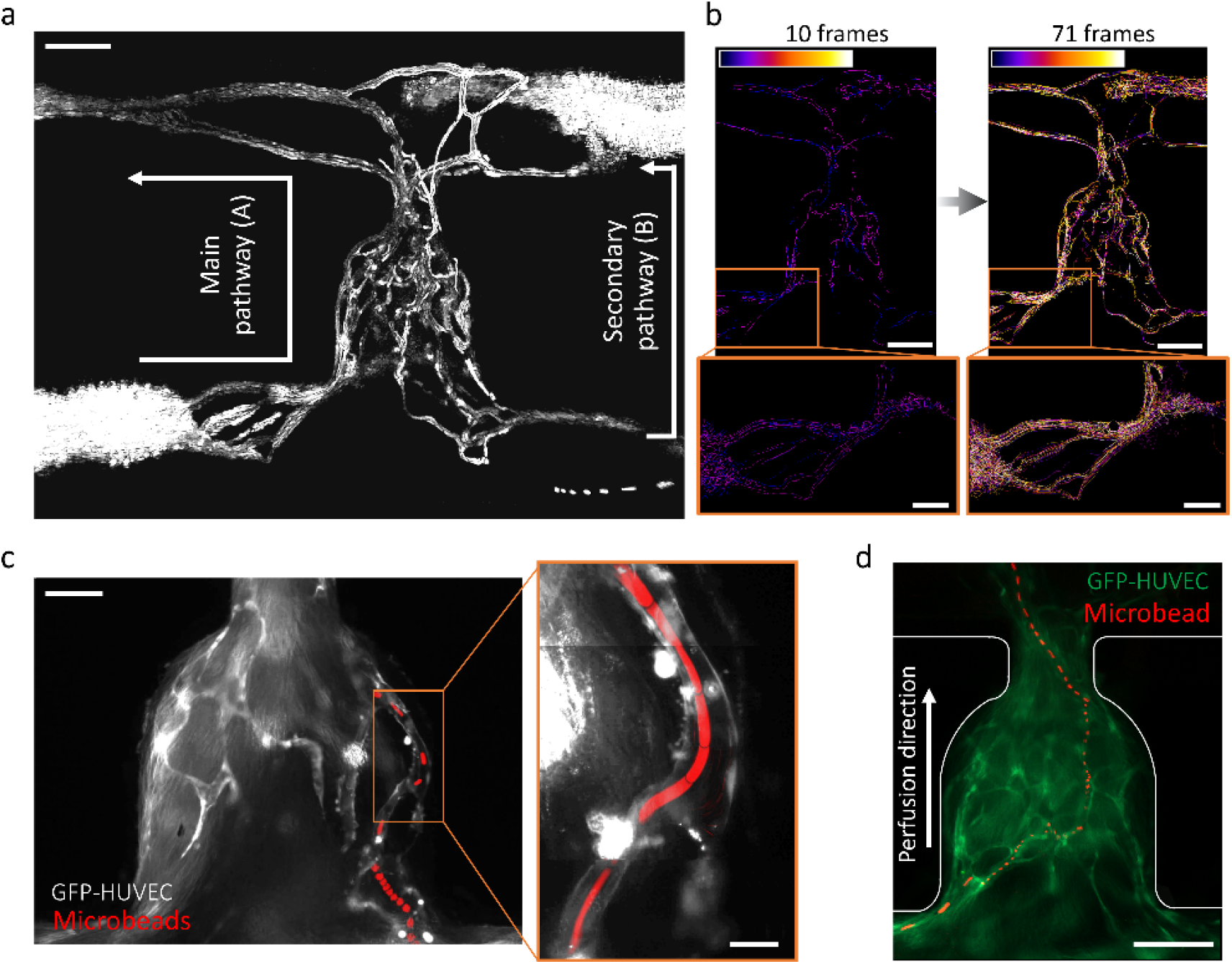
Functionality of the interconnected endothelial networks. **a**, Maximum of intensity projection over a 71 images stack, highlighting the tracks of the microbeads passing through the interconnected network. The stack was obtained after an image subtraction process. See Supplementary Video 1 and Supplementary Fig. 5 for raw movies and details. **b**, Sum of the binarized and frame-color coded images from (**a**) showing time-resolved beads perfusion. See Supplementary Video 2 for details. **c-d**, Projections of maximum intensity over an image stack showing tracking of one individual microbead (red) passing through the endothelial network. The inset shows an assembled projection of three movies taken at the indicated area at higher magnification. For better visualization, the RFP-HUVEC cells are not shown in these images. See Supplementary Video 3 for raw data. Beads were 1 μm (**a**), 4.8 μm and 0.5 μm (**c** and inset), and 3.2 μm (**d**) in diameter. Scale bars, 200 μm (**a**, **b** and **d**), 100 μm (**b** (insets) and **c**) and 20 μm (**c** (inset)).

We next analyzed the distribution of the microbeads within the endothelial network over time; this was done by counting the numbers of pixels in each frame of the binarized and frame-color-coded movies that corresponded to the movements of the microbeads. Interestingly, the microbeads perfused the whole endothelial network without any apparent prioritization of any particular area (Fig. 3b, Supplementary Video 2 and Supplementary Fig. 5). In the microchannels shown in Fig. 3c and d, a smaller number of beads was introduced at a lower flow rate of *Q*_−_ = 0.1 μl/min to visualize the motions of individual microbeads (see also Supplementary Video 3). Superposition of the tracked microbeads (red signals) and the green fluorescent signal coming from the mesenchymal cell aggregate endothelium confirmed perfusion of the network. These results demonstrate that the integrated endothelial networks are functional and readily perfusable.

Lastly, we investigated the physiological relevance of our microfluidic platform. The observed flows in the recorded networks were characterised by tracking fluorescent microbeads as they entered the endothelial network. Depending on the flow rate imposed by the syringe pump, microbeads moving through the microvessels exhibited fluid velocities ranging from *v*_*min*_ = 100 *μ*m/s to *v*_*max*_ = 7500 *μ*m/s. Assuming laminar flow, one can deduct from these values the shear rate in the fluid at the vessel wall, which is given by 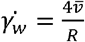, where *R* is the radius of the vessel and 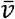 the linear fluid velocity (Supplementary Note 2). Thus, 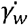 ranged from 27 to 2000 s^−1^. The mean (± s.d.) velocity over the perfused network at *Q*_−,_ and associated shear rate, were 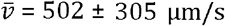 and 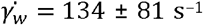, respectively (n = 4 beads individually tracked). The mean (± s.d.) velocity over the perfused network at *Q*_*+*_, and the associated wall shear rate, were 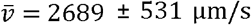 and 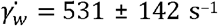, respectively (n = 4 beads individually tracked) (Table 1). The observed range of flux values was consistent with the flow rates detected in the human capillaries, corresponding to velocities of 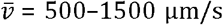 and wall shear rates of 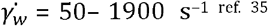. Thus, the observed perfusion flow in our organ-on-chip device resembles physiologic flow observed in human capillaries.

**Table 1.** Summary of different flow parameters measured on the organ-on-chips as compared to *in vivo* physiological flow rates in human capillaries. Data represent mean values ± s.d.

| Experimental flow-rate ($\mu\text{L}/\text{min}$) | Velocity ($\mu\text{m}/\text{s}$) | | Shear rate ($\text{s}^{-1}$) | |
| --- | --- | --- | --- | --- |
|  | On-chip | Physiological | On-chip | Physiological |
| $Q_- = 0.1$ | $v_{\min} = 100$<br>$\bar{v} = 502 \pm 305$ | $v_{\min} = 500$ | $\dot{\gamma}_{w \min} = 26$<br>$\dot{\gamma}_w = 134 \pm 81$ | $\dot{\gamma}_{w \min} = 50$ |
| $Q_+ = 10$ | $\bar{v} = 2689 \pm 531$<br>$v_{\max} = 7500$ | $v_{\max} = 1500$ | $\dot{\gamma}_w = 531 \pm 142$<br>$\dot{\gamma}_{w \max} = 2000$ | $\dot{\gamma}_{w \max} = 1900$ |

### Blood vessel organoids

While significant progress has been made in the development of complex organoids, the convergence of human tissue engineering and microfluidics is needed to address the technical challenges that remain, in particular to vascularize 3D tissues to support extended growth and differentiation^25^. To further show the utility of our device design, we seeded 3D human blood vessel organoids (BVOs) generated from human-induced pluripotent stem cells (hiPSCs) onto our chip. We have previously reported the generation of such organoids^36^ and demonstrated their physiological relevance for modelling diabetic vasculopathy^26^. Slight modifications were made to the BVO differentiation protocol to ensure that the organoids were homogenous and fit into the microfluidic chip and the U-shaped trap, detailed in the Methods section. Importantly, the organoids self-organize into three-dimensional interconnected networks of bona fide capillaries containing an endothelial cell lined lumen, pericyte coverage and a prototypic basal membrane.

Mature blood vessel organoids, GFP-HUVEC cells (6×10^6^ cells/ml) and fibroblasts (2×10^6^ cells/ml) were embedded within the hydrogel and introduced into the microchannels as described above. The HUVEC cells self-organized into endothelial networks arborizing the seeded BVOs after a few days in culture (Fig. 4a). HUVEC networks with hollow lumen structures were observed after on-chip immunostaining on day 13 of culture (Fig. 4b). All the GFP-HUVEC vessels were CD31 positive (CD31^+^) after immunostaining, showing that the microvascular networks can be readily perfused with the anti-CD31 antibody (Fig. 4c). Functionality of the HUVEC networks was also assessed by the perfusion of fluorescent microbeads through the microchannels after 13 days of culture (Fig. 4d, Supplementary Video 4). We determined the reproducibility of our method by quantifying the developed microvascular networks in the U-cup trap area after 10 to 14 days of culture from independent experiments, *i*.*e*. from different batch of organoids, cells, hydrogels and microfluidic chips. The number of junctions, the number of segments, the total segment length and the number of meshes were evaluated using the Angiogenesis Analyzer plugin in ImageJ (Supplementary Fig. 6). No significant difference between the experiments was found (Fig. 4e). Thus, these findings demonstrate the robustness of our microfluidic platform to form stable endothelial networks in a reproducible way, arborizing the trapped BVO.

**Fig. 4.**
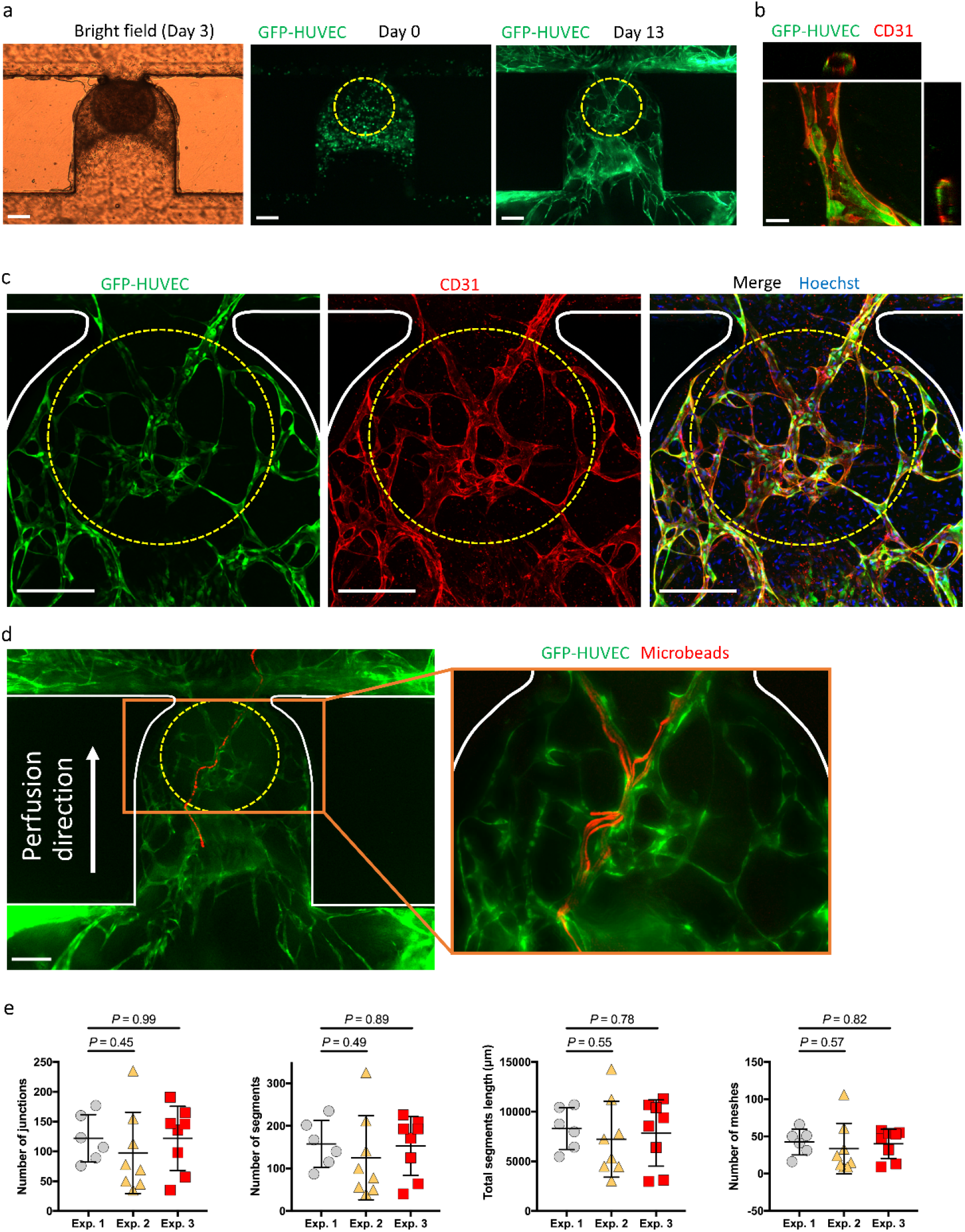
Establishment of perfusable HUVEC endothelial networks around blood vessel organoids. **a**, Time resolved evolution of the cell culture on chip. GFP-HUVEC cells self-organized into an endothelial network surrounding the organoid (circled in yellow from phase contrast images). **b**, Vessel from the endothelial HUVEC network (stained for the endothelial marker CD31) with orthogonal views showing hollow lumen structures. **c**, Confocal z-stack maximum intensity projection of the endothelial network at day 14 after staining of the microfluidic chip for CD31 expression and the nuclear marker Hoechst. **d**, Projection of maximum intensity over an image stack showing the tracking of one individual microbead (red) passing through the endothelial network (green). The inset shows the perfusion of several microbeads at higher magnification. See Supplementary Video 4 for raw movies. **e**, Angiogenesis Analyzer output of the endothelial networks after two weeks of culture for four parameters: (i) numbers of junctions, (ii) number of segments, (iii) total segments length and (iv) number of meshes. Experiments were repeated on n = 22 microchannels from n = 3 different experiments and data are plotted as mean ± s.d. Scale bars, 200 μm (**a**, **c**, **d**) and 20 μm (**b**).

Next, we examined anastomoses between the endothelial networks formed by HUVEC cells and the seeded BVOs. Proper vascularization of organoids remains a major challenge in the field^37^. Having previously shown that BVOs transplanted into the kidney capsule of mice formed functional connections with the host vasculature, we tested whether we could achieve the same *in vitro* with the HUVEC endothelial network. After 10 to 14 days of on-chip culture, the microchannels were fixed and stained *via* a dynamic on-chip protocol where PFA 4%, blocking buffer, primary antibody, secondary antibody and Hoechst solutions were sequentially flowed into the microchannels. Red fluorescent-labelled anti-CD31 antibody staining revealed staining of endothelial cells internal to the BVO vasculature (Fig. 5a). Numerous BVOs vessels (CD31^+^GFP^-^) were observed near GFP-HUVEC vessels, indicating functional perfusion from the HUVEC endothelial bed into the organoids’ vessels (Fig. 5b). Importantly, we were also able to identify several interfaces showing anastomosis between the HUVEC endothelial network and BVO vessels (Fig. 5c and Supplementary Fig. 7). Of note, in regions where the HUVEC network was very dense, the BVOs were concealed, making them difficult to observe. Thus, we plotted the region of interest mean values from z-stacks of green (HUVECs) and red (CD31 staining detecting HUVEC and BVO endothelial cells) channels (z-axis profiles) for 6 different BVOs-on-chip (Fig. 5d). In the analyzed areas, red and green signal curves overlapped in the first images of the z-stacks (close to the bottom of the chip) and diverged when reaching the organoid. This analysis further confirmed that GFP^-^ BVO vessels were perfused and labelled with the anti-CD31 antibody.

**Fig. 5.**
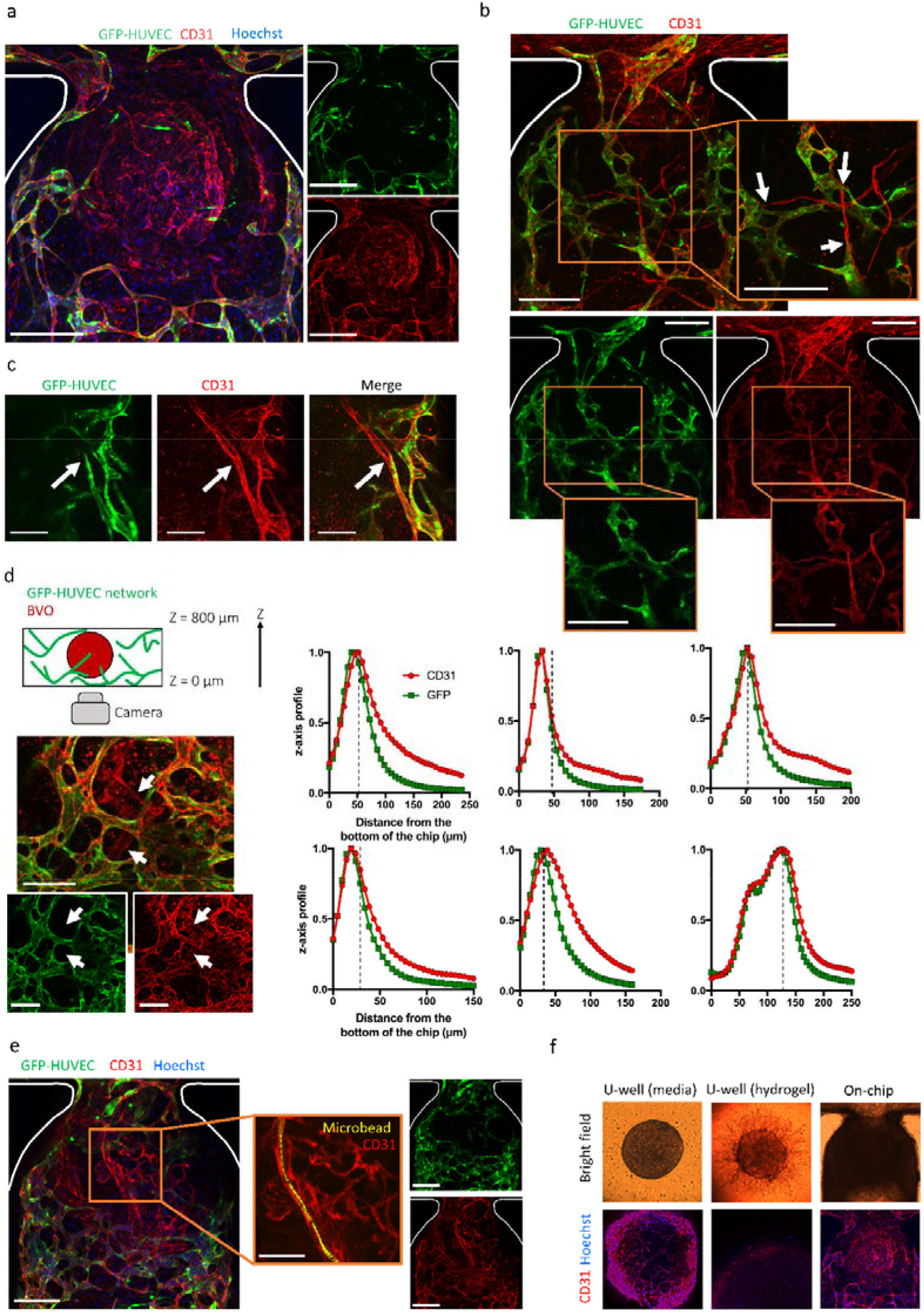
Assessment of functional anastomosis between an HUVEC endothelial bed and stem cell engineered blood vessel organoids. **a-b**, Confocal z-stacks maximum intensity projection at day 10 after staining of the microvessels for CD31 expression (red) and the nuclear marker Hoechst. The BVO vasculature corresponding to CD31^+^GFP^-^ vessels isvisible in the center of the trap. Areas where BVO vessels (CD31^+^GFP^-^) merge with the endothelial HUVEC network (CD31^+^GFP^+^) are indicated with white arrows. **c**, Anastomosis of a GFP^+^ HUVEC vessel with iPS cell-derived blood vessels of the organoid. **d**, Confocal z-stack maximum intensity projection after immunostaining to detect CD31^+^ (red) endothelial cells and detection of GFP^+^ (green) HUVECs (left). The image is representative of n = 6 independent microchannels from n = 3 different experiments for which the z-axis profiles are shown (right). For each graph, a vertical dotted line highlights the divergence of green and red signals, revealing the BVO immunostaining deep inside the microfluidic chip. **e,** Projection of maximum intensity over an image stack showing the tracking of one individual microbead (yellow) passing through the BVO endothelium. Right panels show red and green color channels separately. **f,** Phase contrast images and confocal z-stacks maximum intensity projection for vascular and nuclear markers of BVOs immersed in growth medium (U-well), embedded in hydrogel (U-well) and vascularized on-chip. Scale bars, 200 μm (**a**, **b**, **e** and **f**), 100 μm (**c**, **d** and **e** (inset)).

Finally, we performed microbead perfusions on-chip. Since the beads preferentially flow along the paths of least fluid resistance, the likelihood that they will enter the narrow vasculature of the organoid is low. Nevertheless, with a large number of infused beads, we were able to demonstrate the passage of 2 μm microbeads within large vessels of the organoids (Fig. 5e and Supplementary Fig. 8). Control experiments were performed with BVOs cultured in low-attachment 96 U-wells either immersed in media or embedded in hydrogel. Phase contrast images showed that endothelial sprouts were able to expand in the fibrin hydrogel after a few days of culture. However, immunostaining of BVOs cultured in such conditions was not achievable, showing that neither the antibodies nor the Hoechst molecules were able to diffuse through the hydrogel, contrary to on-chip cultures where the antibodies can reach the BVO *via* the endothelial network and its functional connections with BVO vasculature. Together, these results show that our method is suitable for connecting HUVEC endothelial networks with bona fide blood vessel organoids to organize a perfusable vascular network. This *in vitro* microfluidic system represents a significant step forward in the field of vascularized organoids-on-chip and demonstrates the possibility of infusing different molecules (small-molecule drugs, nucleic acids, antibodies, etc.) or cells into the stem cell-derived blood vessel organoids through the microchannels.

## Discussion

In this paper we present a new microfluidic platform which provides simple and robust solutions to form vascularized organoids-on-chip with precise control over the fluxes generated during short- and long-term perfusion. Through this new approach, we can counteract confining issues often encountered by organ-on-chip technologies, such as the numerous manual operations and lack of automation in spheroid or organoid location and hydrogel handling. First, the device was manufactured using COC, a material readily adapted by industry, avoiding the complications of PDMS systems such as incompatibility with hydrophobic compounds or adsorption of molecules. Second, in most studies, the flow rate is determined by hydrostatic pressure, a simple and cost-efficient liquid actuation principle. However, the optimum hydrostatic pressure difference cannot be maintained for a long time, with the consequence that the culture medium has to be changed regularly^6,38,39^. This can be avoided by using rocker perfusion platforms^14,40^, but this is not representative of *in vivo* flows. In our study, this has been overcome through the use of a syringe pump. Using 20 ml syringes, the microfluidic perfusion can be maintained for two weeks at a flow rate of 1 μl/min without disconnecting the syringes. It should be noted that by using a 10-channel syringe pump, as was done in this study, flows in 10 different channels can be easily monitored in parallel. Third, the organoids and hydrogels were passively placed into a hydrodynamic trap, wherein the cells were subjected to minimal shear stress. These traps can designed in series, thus hosting multiple vascularized organoids (including different tissue organoids) interconnected through endothelial networks.

Cell and hydrogel loading, as well as long-term organoid perfusion with the growth medium, were performed through a single fluidic inlet. This automated method of fluid handling provides the simplicity required for distributing microfluidic devices to industries or hospitals. Finally, the presented microfluidic platform offers robustness through an innovative encapsulation technique, which relies on hydrodynamic and capillary effects that are largely independent of the working pressure. While gel loading is often a delicate step in conventional chip designs found in the literature, the process used in this study is highly reproducible and easy to perform. The loading steps can be performed in seconds, allowing all operations to be completed at room temperature. In summary, this new organ-on-chip platform is very robust and user-friendly, overcoming some of the key hurdles to industrialize organs-on-chips and to provide greater access to user-friendly organoid-on-chip technologies.

Our microfluidic device permits the vascularization and perfusion of a biological 3D tissues. The formation of endothelial networks was found to be highly reproducible. Importantly, we were able to generate physiological fluxes in these networks, with observed velocities ranging from *v* = 100 *μ*m/s to *v* = 7500 *μ*m/s and shear rates ranging from 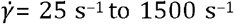. Thus, blood flow velocities and shear rates in human capillaries, ranging from 500 to 1500 *μ*m/s and 100 to 500 s^−1^, respectively, can be readily reached using our microfluidic platform. This contrasts with what has been reported in literature, wherein devices operated by hydrostatic pressure often lead to lower flow rates^22^. Finally, the deposition of the hydrogel on the walls of the microchannels, conditioned by the Landau-Levich-Bretherton effect, offers a robust method of endothelialization of the perfusion channel. This eliminates the need to coat the channels and tilt the chips to adhere cells to the walls, as is often reported^41,42^.

The chip configuration described here resulted in anastomosis between an HUVEC endothelial bed and the endothelium of our blood vessel organoids. These BVOs represent bona fide capillaries with a defined lumen, endothelial lining, pericytes, and formation of a prototypic basal membrane^36^. However, the critical next step to use these human stem cell-derived BVOs to for instance model disease processes was never accomplished so far, namely to generate BVOs which can be perfused. Most importantly, in our current study, we have now accomplished intravascular perfusion of blood vessel organoids, thus making our microfluidic platform the first device to incorporate functional vasculature throughout the microfluidic channel into the embedded organoids. Our platform allows many diverse topics to be studied, such as organoid lifespan enhancement through vascularization, circulation and fate of immune cells in organoids, exposure to drugs, nucleic acids or metabolic stress. The device we have developed also offers the flexibility to vascularize other types of pre-endothelialized organoids, spheroids, tumoroids, or human tissue explants.

## Methods

### Trapping principle and cell seeding

A syringe was used as a reservoir at the entrance of the microfluidic channel to inject three fluid phases in the following order: the hydrogel (1) in which spheroids/organoids and endothelial cells were embedded, the air (2), and the growth medium (3). The microfluidic channel was composed of a serpentine-shaped loop channel with a narrow U-cup-shaped region that functioned as a cell aggregate trap. When the trap was empty, the hydraulic resistance *R*_*1*_ along the trap (Path A) was lower than the one along the loop channel (*R*_*2*,_ Path B). As a result, a spheroid/organoid in the flow was carried into the trap. The loading of different components was done through a syringe pump connected to the outlet of the channel in its withdrawal mode at a flow rate of *Q* = 300 μl/min. A spheroid/organoid was collected from a 96-well plate using a pipette tip and then put in the hydrogel mixture. 50 μl of the hydrogel containing the spheroid/organoid was put into the reservoir at the entrance of the device and introduced in the channel, where the spheroid/organoid was trapped inside the U-cup-shaped microchamber. Air was introduced in the same manner before the hydrogel’s polymerization. The hydrogel was thus pushed to the outlet of the channel. Due to the capillarity, the gel remained in the U-cup site where the spheroid/organoid was trapped, and a thin layer of gel coated the corners of the channel. After the gel’s polymerisation at room temperature for 15 min, growth medium was introduced. Using this set-up, the organoid was affixed in a central microchamber, surrounded by a perfusion channel to allow flow of the growth medium. Finally, the organ-on-a-chip setup was placed inside an incubator at 37°C and 5% CO_2_ (Supplementary Fig. 1).

### Cell culture and generation of spheroids and organoids

Primary human fibroblasts were extracted from skin explants obtained through the elective breast surgery of a healthy young woman following informed consent; this tissue was provided by Walid Rachidi, CEA, Grenoble. GFP- and RFP-labelled. HUVEC cells were purchased from Angio-Proteomie (Boston, MA, US) and cultured in complete EndoGM medium (Angio-Proteomie). Passage 5–7 cells were used for the experiments. Fibroblasts cultured in DMEM, high glucose, GlutaMAX Supplement (Gibco, Grand Island, NY, US), and passage 6–8 cells were used for the experiments. We prepared fibroblasts and HUVEC co-culture, termed mesenchymal cell spheroid model here, in a U-shaped 96-well microplate with an ultra-low attachment surface (Corning, NY, US). Fibroblasts and HUVEC cells were mixed at a ratio of 1:1 (5000 cells per well) in 150 μl of medium. After pre-culturing for 1 day in the microplate, a spheroid was introduced into the device. A mix of CnT-ENDO / CnT-Prime Fibroblast medium (CELLnTEC, Bern, Switzerland) 1% Pen-Strep (PAN Biotech, Aidenbach, Germany) was used for the microfluidic perfusion of the fibroblasts and HUVEC co-culture spheroids. RFP-HUVEC cells were suspended in the hydrogel at a concentration of 6×10^6^ cells per ml.

3D human blood vessel organoids were generated from human induced pluripotent stem cells (hiPSCs) as previously described^36^. In brief, stem cell colonies were harvested using Accutase (SigmaAldrich) to get a single cell suspension. To make sure that the organoids would be of an appropriate size to fit into the 800 μm x 800 μm square profile microchannels of the chip used for BVO experiments, AggreWell™400 (STEMCELL Technologies) plates were used. Each well of the plate contains 1200 microwells with a 400 μm diameter. 600000 single stem cells were seeded per well (500 cells/microwell) in aggregation media with 50 μM Y27632 (STEMCELL technologies). Mesoderm induction and sprouting was induced directly in the AggreWell™400 plates by carefully changing the media with a p1000 pipette making sure to not disturb the cell aggregates in the microwells. For Collagen I-Matrigel embedding, organoids were harvested from the AggreWell™400 plate by vigorously pipetting up and down with a cut p1000 tip close to the bottom of the well. Harvested organoids were embedded in a 12-well plate (approx. 100 organoids/12-well), and subsequently cut out and singled into low attachment 96 U-well plates 4-5 days after embedding as previously described^36^. No alterations were made to any of the BVO differentiation media. The BVOs were maintained in a differentiation medium containing 15% FBS (Gibco), 100 ng/ml VEGF-A and 100 ng/ml FGF-2. The BVOs grew slightly after removal from the AggreWell™400 plate, so 4 days after singling organoids with a diameter of 500-600 μm were selected and added to the microfluidic chip. After adding BVOs to the microfluidic chip, a mix of the differentiation medium with CnT-ENDO and CnT Prime Fibroblast media was used (ratio 1:1:1).

### Hydrogel preparation

A fibrin-hydrogel made of 6.6 mg/ml fibrinogen, 0.15 TIU/ml aprotinin, 2.5 mM CaCl_2_, and 1 U/ml thrombin prepared in HEPES-buffered saline (Sigma–Aldrich, Taufkirchen, Germany) was used in all experiments. After adding the thrombin (Sigma–Aldrich, St. Louis, MO, USA) into the mixture, all the procedures were quickly performed to avoid premature gelation.

### Device fabrication

Computer-aided design (CAD) files of the chip were generated using Solidworks 2018 (Dassault Systèmes, Vélizy-Villacoublay, France). The microfluidic chips were made of Cyclic Olefin Copolymer (COC) because of its low autofluorescence, strong chemical resistance, and low drug absorption. Microfluidic patterns were directly machined on a COC sheet (TOPAS, USA), using high-precision milling (DATRON M7HP equipment). The chip (84×54 mm2) contained 10 identical microfluidic channels. The square-profiled channels were 400 μm x 400 μm for the experiments with mesenchymal spheroids, and 800 μm x 800 μm for experiments with blood vessel organoids. The microfluidic channels were sealed with a MicroAmp optical adhesive film (Applied Biosystems, Foster City, USA).

### Cell imaging

The microfluidic chip was imaged using an inverted Olympus IX50 microscope every day for the period of the experiment. Images were taken in bright field, green fluorescence, and red fluorescence channels, with 4x, 0.1 N.A and 10x, 0.3 N.A objectives, equipped with a CCD camera interfaced with Point Grey software. Images showing anastomosis were taken using an inverted microscope (Zeiss Observer Z1) with a 20x, 0.4 N.A objective equipped with an Axiocam 503 mono CCD camera and ApoTome technology (optical sectioning using structured illumination). Images of the network shown in Fig. 2d were taken using a confocal spinning disk system, consisting of an EclipseTi-E Nikon inverted microscope, equipped with a CSUX1-A1 Yokogawa confocal head, an Evolve EMCCD camera (Roper Scientific, Princeton Instruments), and a 20x, 0.75 N.A. objective, interfaced with MetaMorph software (Universal Imaging). Images of the vascular networks and blood vessel organoids were taken using a confocal microscope, Zeiss LSM800, with 10x, 0.3 N.A and 63x, 1.4 N.A (for hollow lumen) objectives. The flow in vascular networks was assessed in the second week of culture by loading polystyrene fluorescent microbeads (Thermo Fisher Scientific Fluoro-Max Fluorescent Beads) into the serpentine channel. Images were captured at 15 Hz using the inverted Olympus IX50 microscope described above. Microbeads were tracked in perfused tissues from separate microfluidic channels using ImageJ (National Institute of Health, New York, NY, USA).

### Light sheet fluorescence microscopy

A homemade light sheet fluorescence microscope was used in this project, which we adapted to image biological samples inside microfluidic chambers without interfering with the normal function of the chip. The light sheet was generated with a 488 nm Ar-laser, focused by a 100 mm focal length cylindrical lens. The fluorescence signal generated at the illuminated plane was collected by a long working distance, with the objective (Mitutoyo M Plan APO SL 20X, 0.28 N.A.) placed at 90° to the excitation path. The sample plane was at 45° from both of these paths. A tube lens was associated to the objective to form the image of the fluorescent structure onto a high-sensitive sCMOS camera (Hamamatsu HPF6 ORCA FLASH 4.0 V3) with a magnification factor of 12. To filter out the laser excitation, a high pass (cut-off wavelength of 490 nm) interference filter was used. The sample was mounted onto a custom-designed holder attached to computer-controlled xz linear translational stages. In this configuration, the microfluidic chip was kept horizontal, and the thinner lateral part of the light sheet was positioned at the surface of the gel. The light sheet illuminated the sample in the direction perpendicular to that of the microfluidic channel (Supplementary Fig. 4).

### On-chip immunofluorescent staining

For immunofluorescent staining, the cells and organoids were fixed by flowing 4% paraformaldehyde (Sigma-Aldrich, Taufkirchen, Germany) for 1 h at room temperature through the microchannel, and subsequently blocked with 3% FBS, 1% BSA, 0.5% Triton-X-100 and 0.5% Tween for 2 h at room temperature. Primary antibodies (CD31, Rabbit, Abcam clone EP3095 ab134168) were diluted at 1:300 in blocking buffer and flowed overnight into the microfluidic chip at 4°C. After a 30 min wash in PBST (0.05% Tween), Cy3-conjugated secondary antibodies (Donkey Anti-Rabbit, Jackson ImmunoResearch Inc.) were flowed into the microchannels at 1:300 in blocking buffer for 2 h at room temperature. After a 30 min wash in PBST, nuclear counter-staining using Hoechst was carried out according to a routine protocol.

### Endothelial networks analysis

Confocal z-stacks of the microchannels in various culture conditions were taken. These stacks were then flattened in ImageJ to a 2D maximum intensity projection and analysed using the Angiogenesis Analyzer plugin with default settings^43^. Three metric parameters were selected for this study, namely the total length of the vascular network, the number of junctions, and the mesh index corresponding to the mean distance separating two master junctions of the trees in the analysed area.

### Statistical analysis

Results are shown as mean ±⍰s.d. as indicated in the Figure legends. Two-way ANOVA with Šidák multiple comparisons tests and unpaired t-test (two-tailed) for comparisons were conducted using GraphPad Prism 9 (GraphPad Software Inc., San Diego, CA, USA). Exact *P* values are provided in the Figures.

## Code availability

Associated code for image analysis is available at: https://github.com/ClementQuintard/Vascularized-Organoids

## Acknowledgments

We thank N.Verplanck and F.Boizot for the manufacturing of microfluidic chips, and X.Mermet for his help with light sheet microscopy experiments. This work was supported by the CEA “OOC inflexion” and received funding from GRAL, a programme from the Chemistry Biology Health (CBH) Graduate School of University Grenoble Alpes (ANR-17-EURE-0003). G.J. and J.M.P. were funded by the Vienna Science and Technology Fund (WWTF) through project COV20-002. J.M.P was further funded by the Austrian Federal Ministry of Education, Science and Research, the Austrian Academy of Sciences and the City of Vienna and grants from the Austrian Science Fund (FWF) Wittgenstein award (Z 271-B19), the T. von Zastrow foundation, the Innovative Medicines Initiative 2 Joint Undertaking (JU) under grant agreement No 101005026, and the Canada 150 Research Chairs Program F18-01336, the Canadian Institutes of Health Research COVID-19 grants F20-02343 and F20-02015, an Allen Distinguished Investigators (ADIs) award, and the Leducq foundation (ReVAMP).

## Contributions

J.M.P., J-L.A., F.N., Y.F. and X.G. conceived and supervised the project; C.Q., C.L. and A.P. performed the experiments; G.J., N.W., A.L. and A.H. generated the blood vessel organoids; C.B., P.B. and C.Q. performed the experiments using light sheet microscope and analyzed the images; C.Q., X.G. and J.M.P wrote the manuscript with inputs and comments from all coauthors. All authors discussed the data and agreed on the final manuscript.

## Supplementary Information

**Supplementary Fig. 1.**
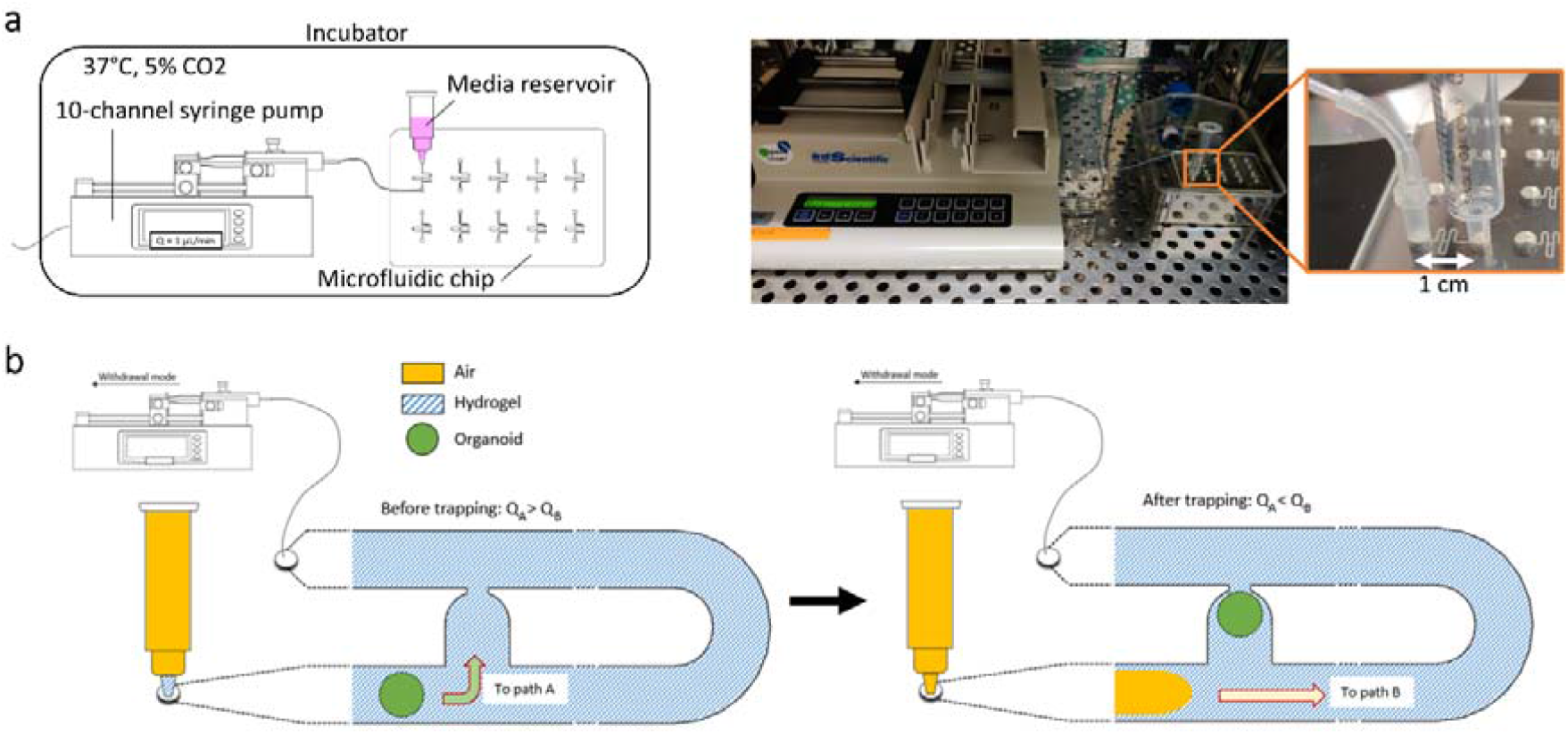
Experimental set-up for loading and continuous perfusion of organoids-on-chip. **a**, Sketch and photograph of the microfluidic platform. Using a 10-channel syringe-pump, the flow can be monitored in the 10 microchannels of the chip. For a better visualization, only one microchannel was connected to the syringe pump on this photograph. **b**, Schematic diagram showing the trapping and encapsulation of an organoid with associated flow rates along paths A and B (Q_A_ and Q_B_, respectively). See Supplementary Note 1 for details regarding essential flow rate criterions for proper function of the encapsulation and continued perfusion mechanisms.

**Supplementary Fig. 2.**
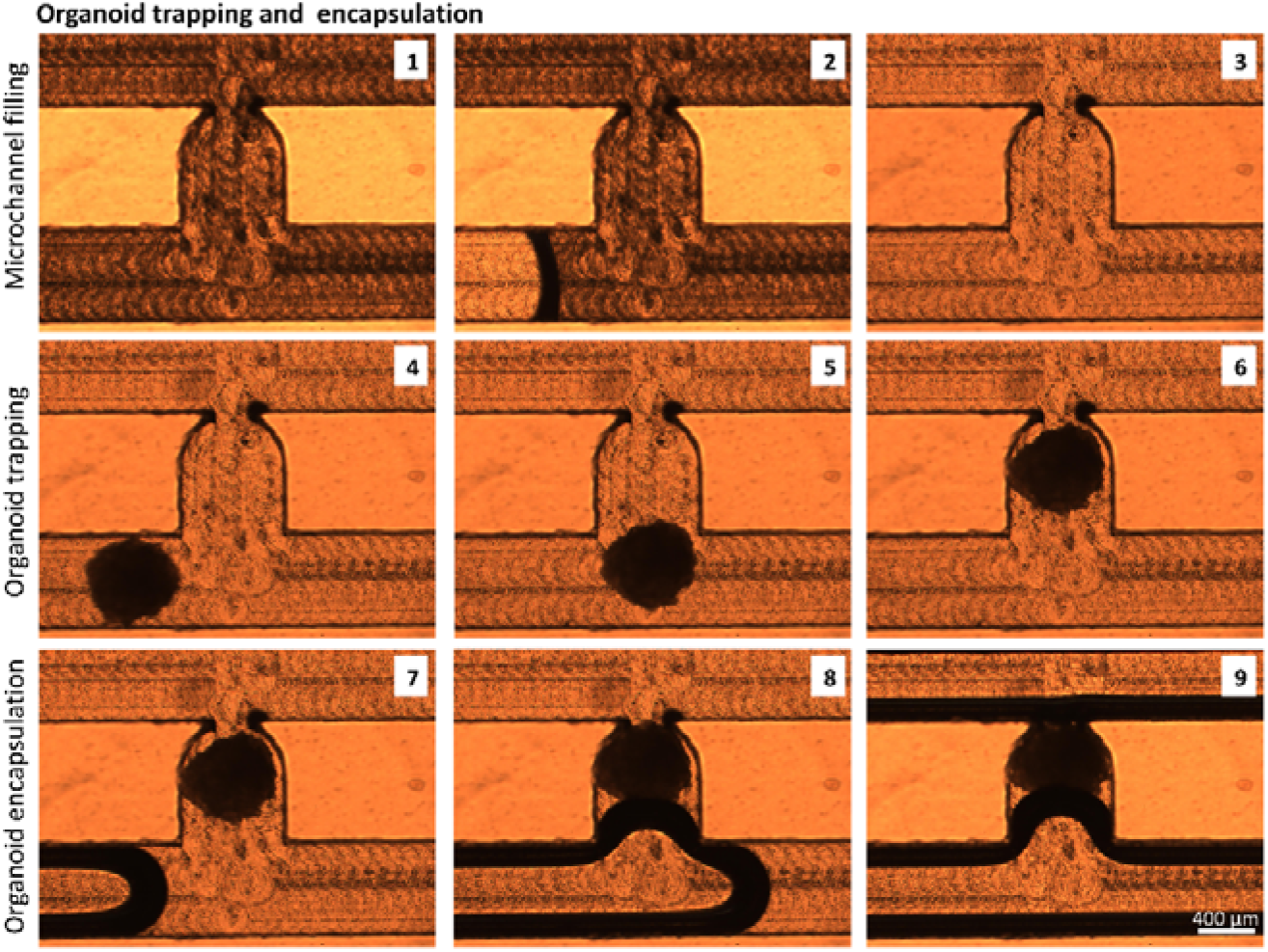
Experimental validation of organoid trapping and encapsulation. Stills from a movie of the experimental trapping of one organoid was recorded at 15 fps using an inverted microscope (bright field). First, the hydrogel fills the microchannel (microchannel filling, steps 1-3). Then, the organoid is trapped in the U-cup shaped area (organoid trapping, steps 4-6). Last, the air pushes the hydrogel towards the outlet of the microchannel (organoid encapsulation, steps 7-9). The hydrogel-air interface can be seen in black due to the difference in the refractive index.

**Supplementary Fig. 3.**
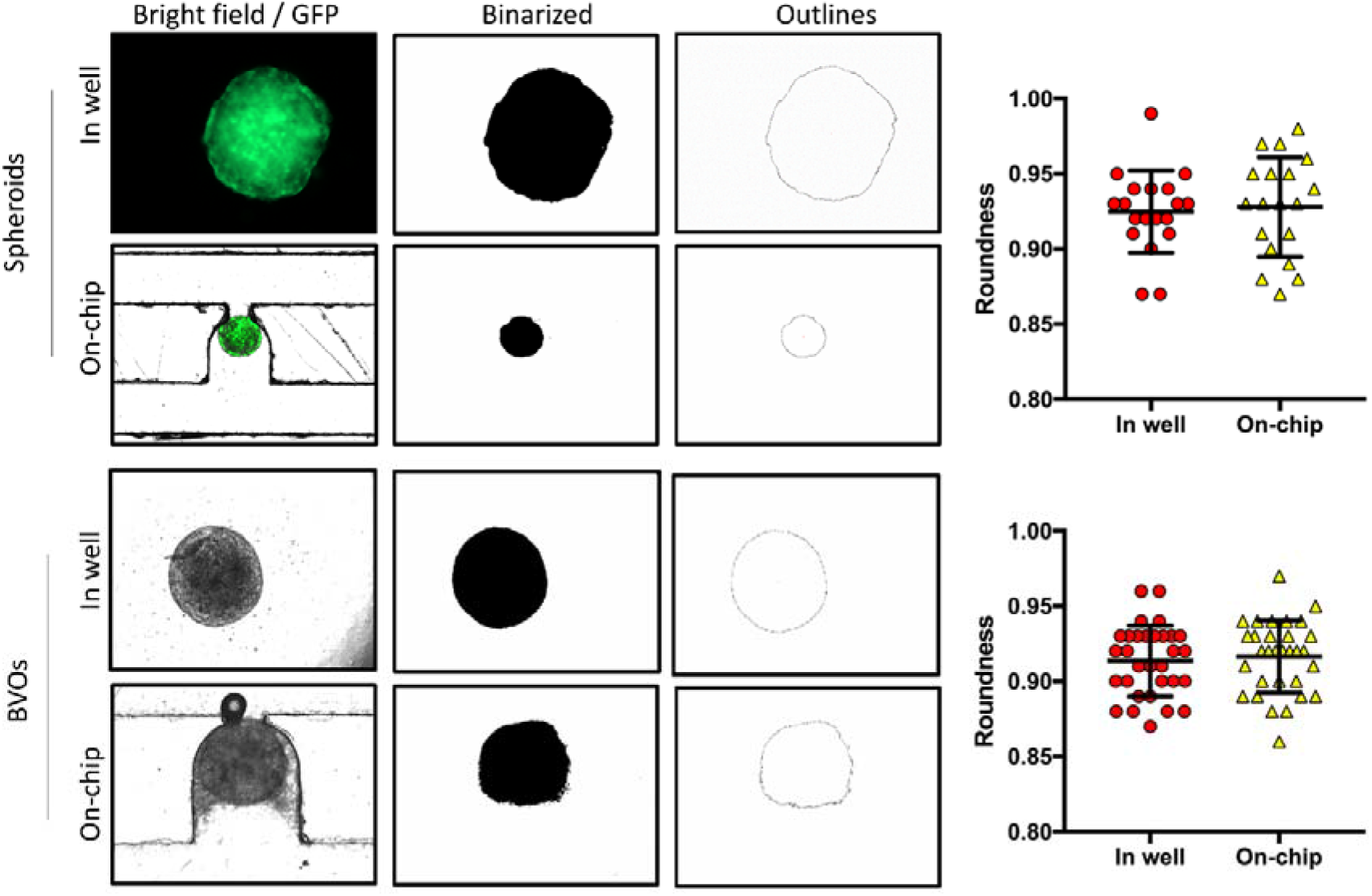
Spheroids and organoids maintained their spherical shape after loading on-chip.. Evaluation of the roundness of spheroids (n = 19) and organoids (n = 31) before and after on-chip encapsulation. The spheroids and organoids were selected according to their spherical shape (roundness above 0.8. Spheroids (n = 19): roundness 0.92 ± 0.03. Organoids (n = 31): roundness 0.92 ± 0.02). They were precisely positioned in the trap site without any apparent morphological alteration, with a roundness maintained above 0.8 (Spheroids (n = 19): roundness 0.93 ± 0.03. Organoids (n = 31): roundness 0.92 ± 0.02). Data represent mean ± s.d.

**Supplementary Fig. 4.**
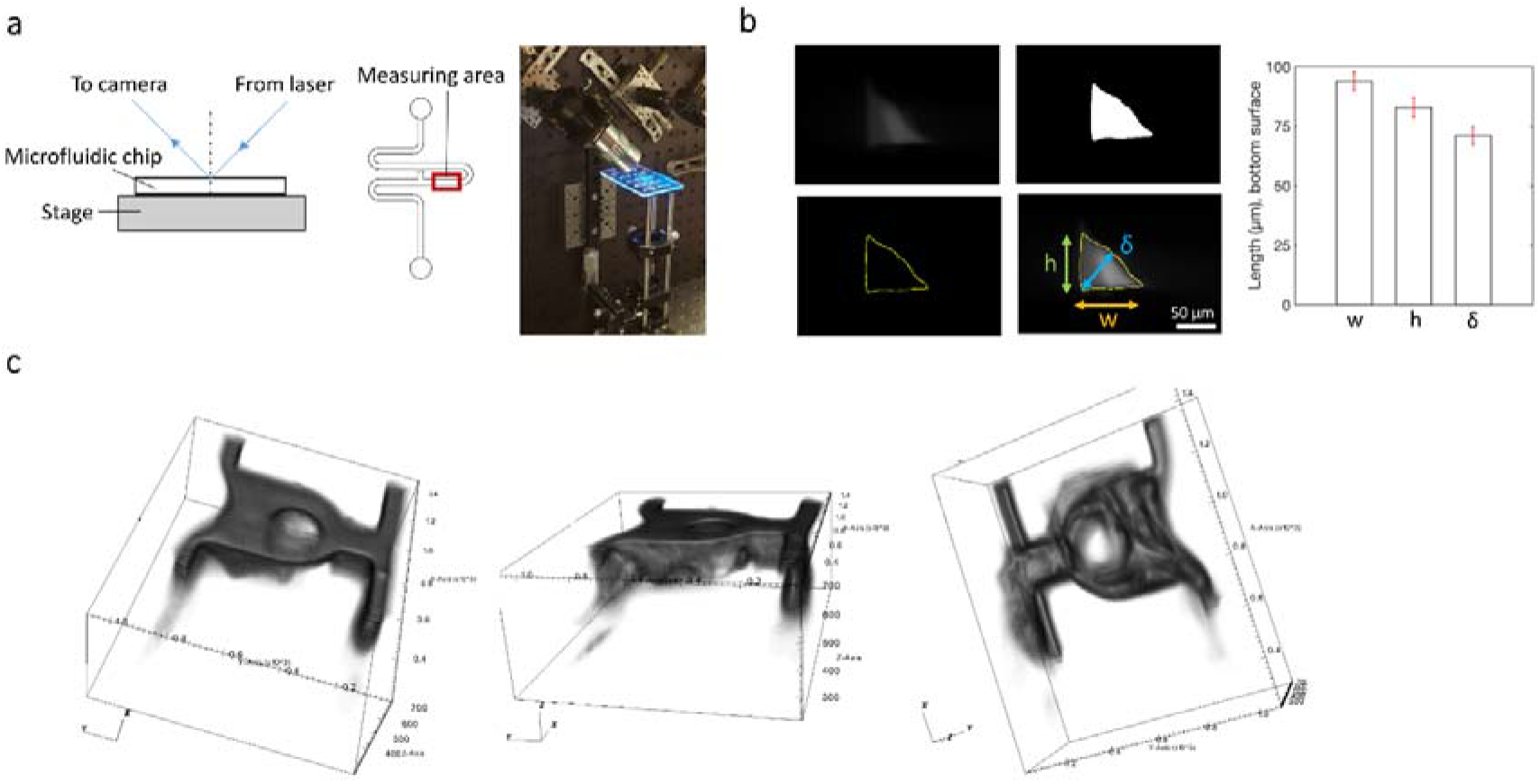
Set-up of experiments using an in-house light sheet microscope. **a**, Sketch and photograph of the optical set-up for light sheet imaging of the chip. A stack of 100 images was acquired in the measuring area to quantify fluorescent hydrogel deposition with a step size of 10 microns along the microfluidic channel using an automated stage. **b**, Images of the hydrogel deposition within one microchannel corner showing raw image, Otsu-based segmentation mask image, segmentation contouring and superposition of raw image and mask. The dimensions of the remaining hydrogel corners in a cross-section of the channel, namely the height (h), the width (w), and the diagonal, Feret’s diameter (δ), were collected. The dimensions represent mean ± s.d. and correspond to an average made over 20 images of the bottom surface, acquired at diverse locations along the microfluidic channel. **c**, Representative 3D views of the fluorescent gel distribution within the microfluidic U-cup trap. The lighter part in the center of the trap corresponds to an air bubble trapped within the gel. A python script was used to re-slice the stack of images collected at 45° to a file containing a 3D intensity matrix. The 3D representations of the trap volume were realized using the VisIt software. Exposure time: 40 ms. laser power: 3 mW.

**Supplementary Fig. 5.**
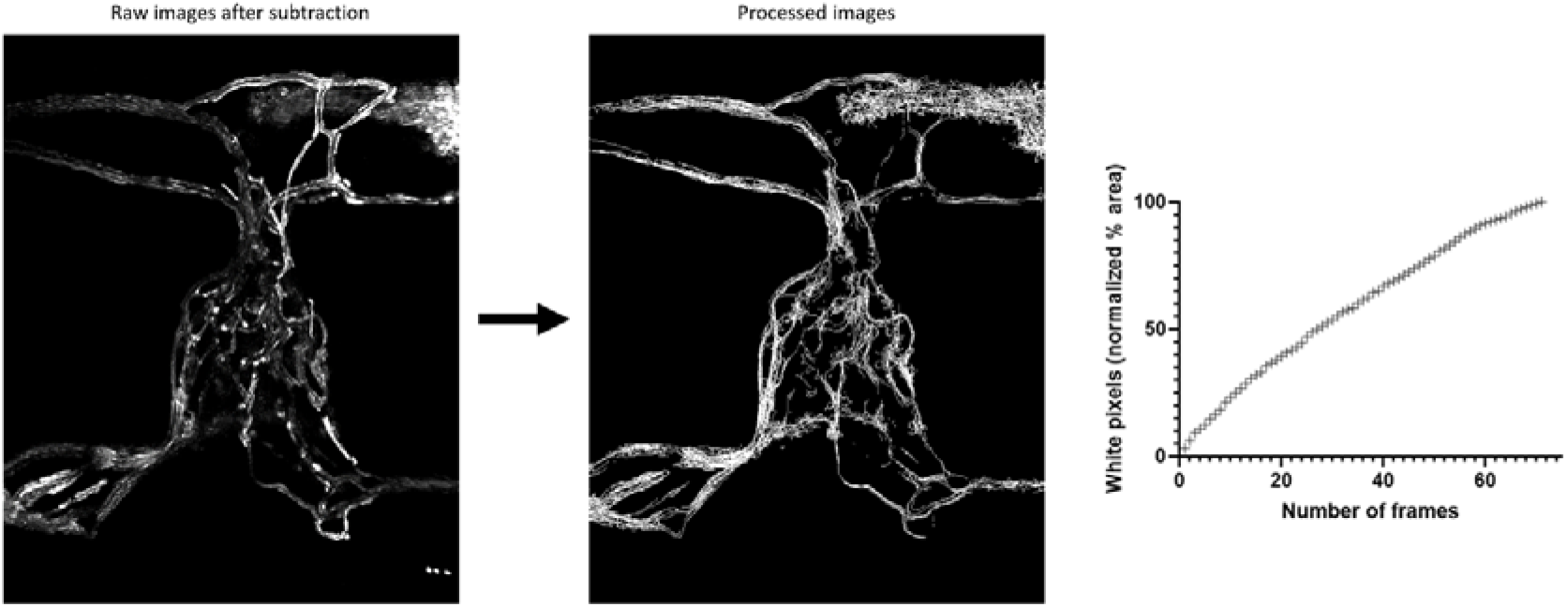
Image processing of microbeads flowing through an endothelial network. Movies of fluorescent microbeads flowing through mesenchymal vascular network were taken at 15 fps using an inverted microscope. Only the traces of the microbeads have been kept using a subtraction script on ImageJ. The images were then process using the Skeletonize and Binarize features in ImageJ, and superimposed frame by frame. The number of white pixels, corresponding to microbeads movements, steadily increased during the experiment, demonstrating that the microbeads do not prioritize any particular vessel of the microvascular network.

**Supplementary Fig. 6.**
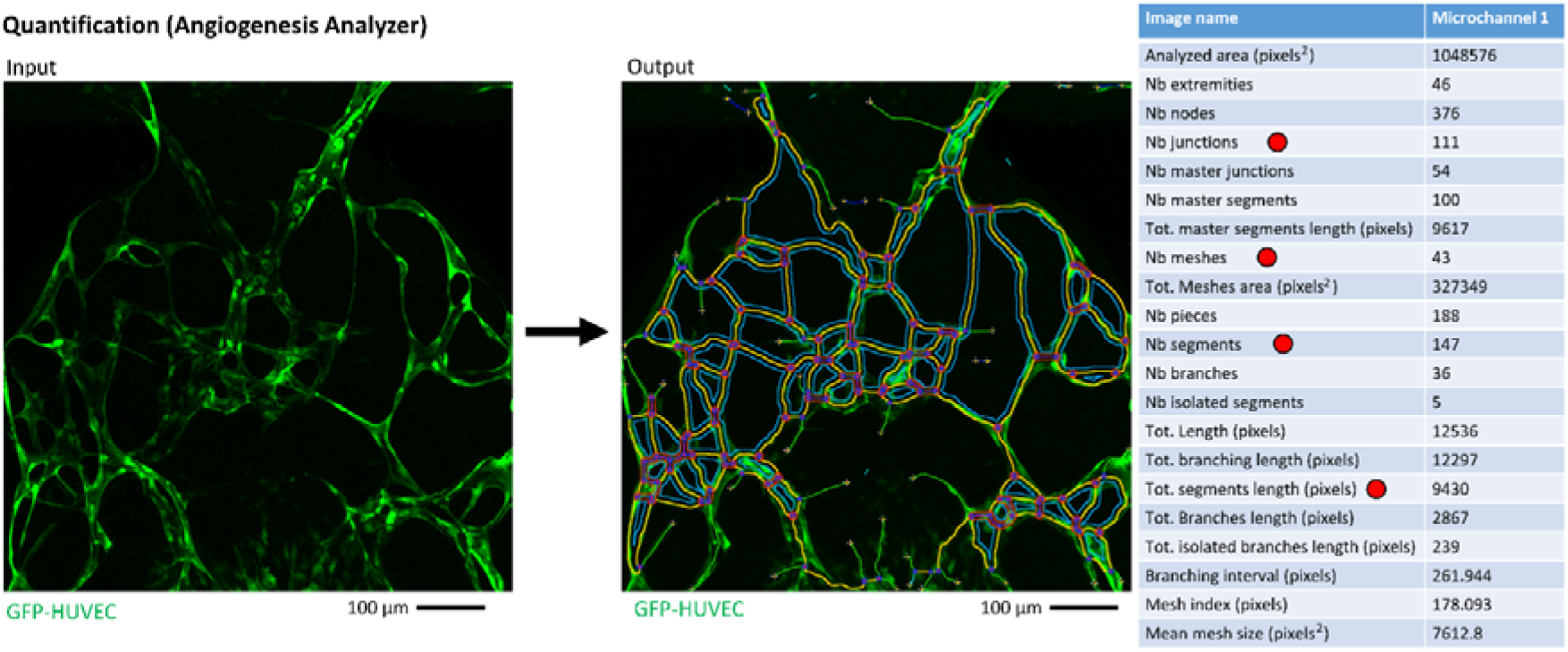
Quantification of endothelial network parameters using the Angiogenesis Analyzer plugin. Confocal z-stacks of the microchannels were taken in the various culture conditions. The z-stacks were taken at the limit of the confocal depth within each sample, nearly 200 μm per sample, which represents approximately the same volume per endothelial network analyzed. Those z-stacks were then flattened in ImageJ to a 2D maximum-intensity projection (as required by Angiogenesis Analyzer). The default settings of the Angiogenesis Analyzer plugin were used. Four different parameters (red dots) representative of the network morphology were used for comparisons.

**Supplementary Fig. 7.**
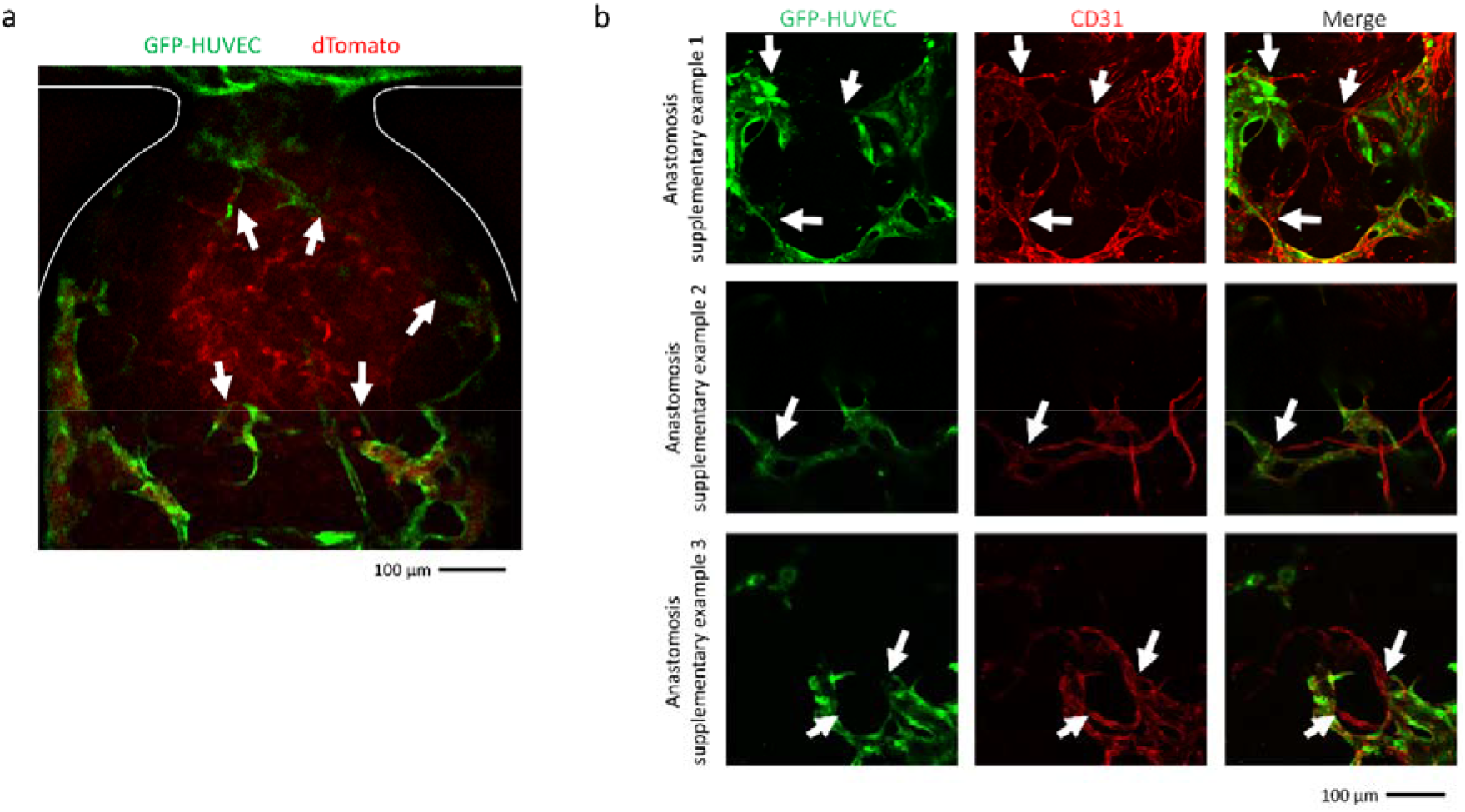
Anastomosis between GFP-HUVEC endothelial networks and blood vessel organoids. **a,** To explore whether connections were able to form between the HUVEC (GFP green) endothelial bed and the trapped organoid, we used blood vessel organoids (BVO) in which the endothelial lineage was fluorescently labelled with dTomato (CDH5-dTomato). Arrows point at anastomoses. **b,** Higher magnification confocal images of a z-plane at the organoid/HUVEC interface showing connections between the GFP-HUVEC vessels and the organoid vasculature (white arrows). Observations were repeated on n = 3 microchannels with similar results.

**Supplementary Fig. 8.**
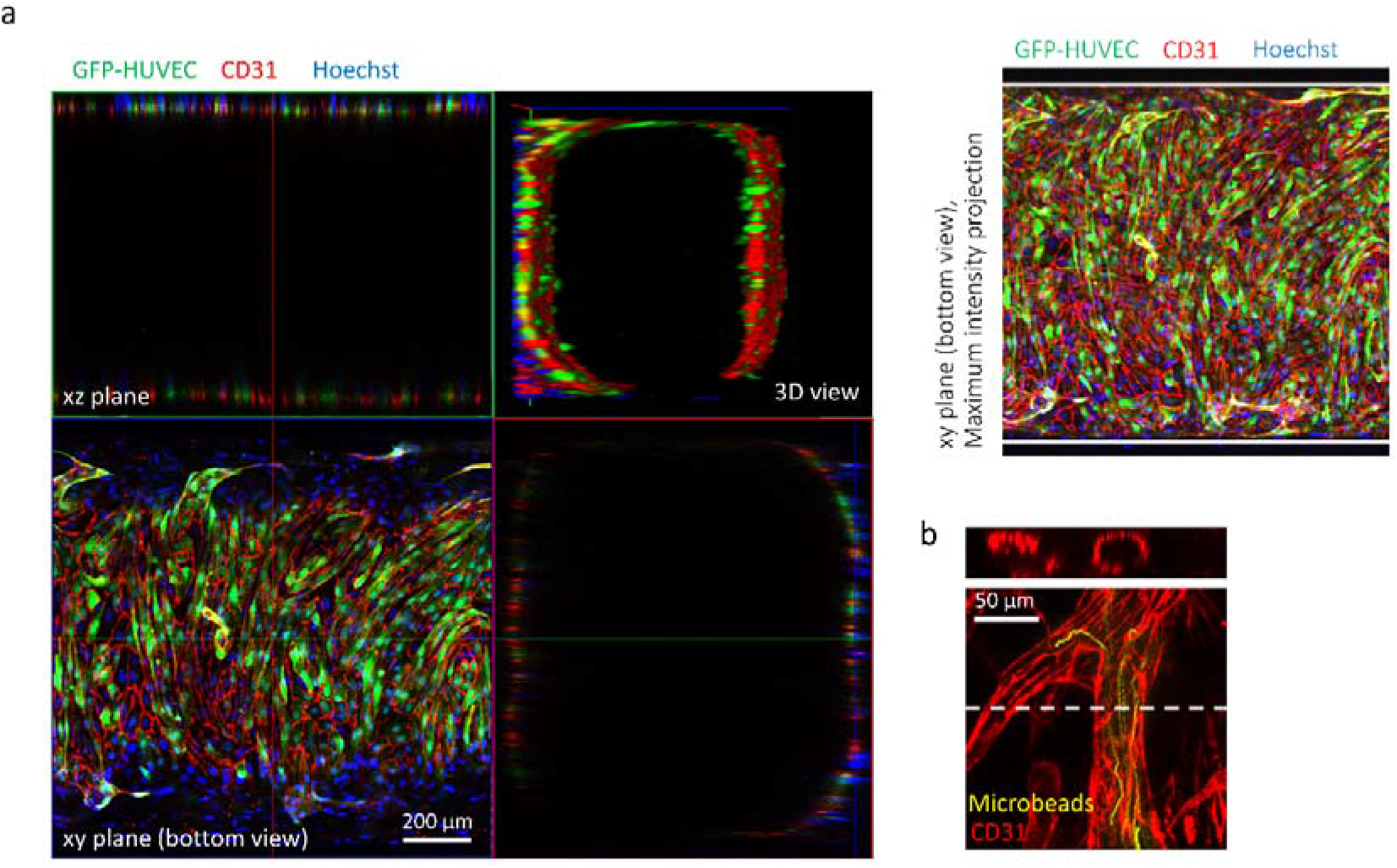
Establishment of a functional vascular tree. **a,** Confocal imaging of the main serpentine channel near the U-cup trap (bottom, orthogonal and 3D views): the serpentine geometry, combined with the loading process presented, allows for the formation of a vascular tree pattern. Due to the hydrogel deposition on the corners of the main serpentine microchannel, the embedded cells rapidly coat the walls of the channel, thus establishing a large vessel in which the growth medium flowed freely for continuous perfusion, branching into smaller vessels of the GFP-HUVEC network and the BVO vasculature shown in **(b). b**, Confocal imaging of an organoid vessel (intensity maximum projection and orthogonal views) showing a hollow lumen structure and perfusion by microbeads.

### Supplementary Note 1: Organoid trapping and encapsulation (theory supplement)

When the spheroid arrives at the trapping unit, it has two options (Supplementary Fig. 9a). Either bypassing the trap via path B, in which case the spheroid is not trapped and heads for the exit. Or the shortcut through path A, in which case the spheroid is trapped in the appropriate area. For the principle of hydrodynamic trapping to work, the spheroid must prefer path A; the microfluidic circuit must therefore globally respect certain geometrical constraints which will be detailed below.

Consider the points *X* and *Y* upstream and downstream of the bypass respectively. Let *ΔP*_*B*_ be the pressure difference between *X* and *Y* passing through the loop (path B) of length *L*_*B*_, and *ΔP*_*A*_ the pressure difference passing through the bypass (path A), whose restriction is of length *L*_*A*_. These pressure differences do not depend on the path followed and are the same.

In the Stokes regime, the pressure differences that occur are linear functions of the liquid flow. Microchannels have rectangular cross sections that induce regular pressure drops of the shape^1^:

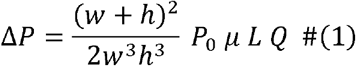

Where *μ* is the dynamic viscosity of the liquid, *w, L* and *h* the width, length and height respectively of the considered channel. *P*_*0*_ is a Poiseuille number characteristic of the cross-sectional geometry of the considered rectangular channel which can be expressed as a function of 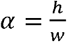 the aspect ratio^2^:

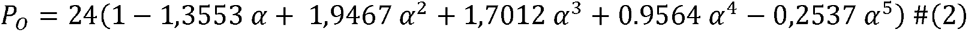

In the case of path B, the channel cross-section is square, hence *α*_*B*_ = 1 and 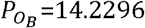, and the pressure difference is written (neglecting the additional pressure drop associated with the bend):

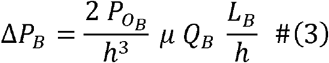

The pressure difference *ΔP*_*A*_ along path *A* will be considered as essentially resulting from the restriction. It is composed of a regular component, due to the friction of the flow along the lateral walls of the restriction of length *L*_*A*_, and a singular component related to the change in cross-section of the flow at the entrance and exit of the restriction (contraction / expansion). Concerning the regular component, by replacing *α* by 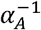 in (2) for the expression of 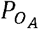 (as gravity plays no role at these scales), one can write the expression for the regular pressure difference associated with path A :

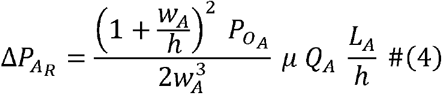

The singular pressure difference can be written as follows^3^:

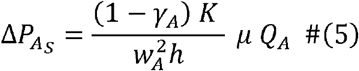

Where we introduced 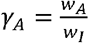 the contraction ratio of path A and *K*, a coefficient that reflects the only effect of the aspect ratio:

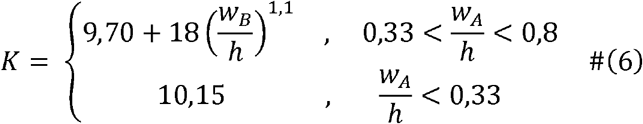

Since the pressure difference between points *X* and *Y* does not depend on the path followed, we have 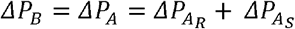 which translates to:

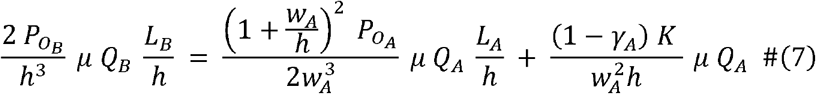

Thus :

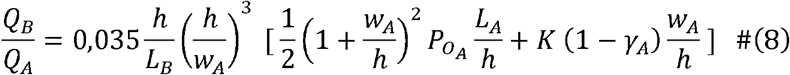

Of note, this final expression does not contain any fluid velocity term, so the hydrodynamic trapping principle works for all velocities in the laminar regime. A spheroid will be trapped if *Q*_*A*_>*Q*_*B*_, so the criterion for a functioning trap is 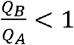.

### Supplementary Note 2: Shear rate at the vessel wall derivation (theory supplement)

Assuming a cylindrical blood vessel of radius *R* along a z-axis, the velocity profile is given by the Poiseuille equation: 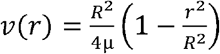. The flow rate *Q* can be expressed as *Q* = ∫*v*(*r*)*dA* with *dA =* 2*πrdr*. Thus, 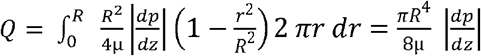. Thus, the average velocity defined by 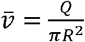 can be expressed as 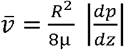. Finally, one can find the shear rate at the blood vessel wall 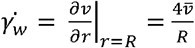.

## References

1. Auger, F. A., Gibot, L. & Lacroix, D. The Pivotal Role of Vascularization in Tissue Engineering. Annu. Rev. Biomed. Eng. 15, 177–200 (2013).

2. van Duinen, V., Trietsch, S. J., Joore, J., Vulto, P. & Hankemeier, T. Microfluidic 3D cell culture: from tools to tissue models. Curr. Opin. Biotechnol. 35, 118–126 (2015).

3. Liu, H. et al. Advances in Hydrogels in Organoids and Organs-on-a-Chip. Adv. Mater. 31, 1902042 (2019).

4. Whisler, J. A., Chen, M. B. & Kamm, R. D. Control of Perfusable Microvascular Network Morphology Using a Multiculture Microfluidic System. Tissue Eng. Part C Methods 20, 543–552 (2012).

5. Kim, S., Lee, H., Chung, M. & Jeon, N. L. Engineering of functional, perfusable 3D microvascular networks on a chip. Lab. Chip 13, 1489–1500 (2013).

6. Alonzo, L. F., Moya, M. L., Shirure, V. S. & George, S. C. Microfluidic device to control interstitial flow-mediated homotypic and heterotypic cellular communication. Lab. Chip 15, 3521–3529 (2015).

7. Nashimoto, Y. et al. Vascularized cancer on a chip: The effect of perfusion on growth and drug delivery of tumor spheroid. Biomaterials 229, 119547 (2020).

8. Paek, J. et al. Microphysiological Engineering of Self-Assembled and Perfusable Microvascular Beds for the Production of Vascularized Three-Dimensional Human Microtissues. ACS Nano 13, 7627–7643 (2019).

9. Shirure, V. S. et al. Tumor-on-a-chip platform to investigate progression and drug sensitivity in cell lines and patient-derived organoids. Lab. Chip 18, 3687–3702 (2018).

10. Sobrino, A. et al. 3D microtumors in vitro supported by perfused vascular networks. Sci. Rep. 6, 1–11 (2016).

11. Rambøl, M. H., Han, E. & Niklason, L. E. Microvessel Network Formation and Interactions with Pancreatic Islets in Three-Dimensional Chip Cultures. Tissue Eng. Part A (2019) doi:10.1089/ten.tea.2019.0186.

12. Homan, K. A. et al. Flow-enhanced vascularization and maturation of kidney organoids in vitro. Nat. Methods 16, 255–262 (2019).

13. Cakir, B. et al. Engineering of human brain organoids with a functional vascular-like system. Nat. Methods 16, 1169–1175 (2019).

14. van Duinen, V. et al. Perfused 3D angiogenic sprouting in a high-throughput in vitro platform. Angiogenesis 22, 157–165 (2019).

15. Campisi, M. et al. 3D self-organized microvascular model of the human blood-brain barrier with endothelial cells, pericytes and astrocytes. Biomaterials 180, 117–129 (2018).

16. Sano, E. et al. Engineering of vascularized 3D cell constructs to model cellular interactions through a vascular network. Biomicrofluidics 12, 042204 (2018).

17. Moya, M. L., Hsu, Y.-H., Lee, A. P., Hughes, C. C. W. & George, S. C. In Vitro Perfused Human Capillary Networks. Tissue Eng. Part C Methods 19, 730–737 (2013).

18. Wang, X. et al. A hydrostatic pressure-driven passive micropump enhanced with siphon-based autofill function. Lab. Chip 18, 2167–2177 (2018).

19. Haase, K., Gillrie, M. R., Hajal, C. & Kamm, R. D. Pericytes Contribute to Dysfunction in a Human 3D Model of Placental Microvasculature through VEGF-Ang-Tie2 Signaling. Adv. Sci. 6, 1900878 (2019).

20. Nashimoto, Y. et al. Integrating perfusable vascular networks with a three-dimensional tissue in a microfluidic device. Integr. Biol. 9, 506–518 (2017).

21. van Duinen, V. et al. 96 perfusable blood vessels to study vascular permeability in vitro. Sci. Rep. 7, 1–11 (2017).

22. Phan, D. T. T. et al. A vascularized and perfused organ-on-a-chip platform for large-scale drug screening applications. Lab. Chip 17, 511–520 (2017).

23. Wang, X., Sun, Q. & Pei, J. Microfluidic-Based 3D Engineered Microvascular Networks and Their Applications in Vascularized Microtumor Models. Micromachines 9, 493 (2018).

24. Zhang, B., Korolj, A., Lai, B. F. L. & Radisic, M. Advances in organ-on-a-chip engineering. Nat. Rev. Mater. 3, 257–278 (2018).

25. Garreta, E. et al. Rethinking organoid technology through bioengineering. Nat. Mater. 1–11 (2020) doi:10.1038/s41563-020-00804-4.

26. Wimmer, R. A. et al. Human blood vessel organoids as a model of diabetic vasculopathy. Nature 565, 505–510 (2019).

27. Jeon, J. S., Chung, S., Kamm, R. D. & Charest, J. L. Hot embossing for fabrication of a microfluidic 3D cell culture platform. Biomed. Microdevices 13, 325–333 (2011).

28. Tan, W.-H. & Takeuchi, S. A trap-and-release integrated microfluidic system for dynamic microarray applications. Proc. Natl. Acad. Sci. 104, 1146–1151 (2007).

29. Tan, W.-H. & Takeuchi, S. Dynamic microarray system with gentle retrieval mechanism for cell-encapsulating hydrogel beads. Lab Chip 8, 259–266 (2008).

30. Zbinden, A. et al. Non-invasive marker-independent high content analysis of a microphysiological human pancreas-on-a-chip model. Matrix Biol. 85–86, 205–220 (2020).

31. Silva, P. N., Green, B. J., Altamentova, S. M. & Rocheleau, J. V. A microfluidic device designed to induce media flow throughout pancreatic islets while limiting shear-induced damage. Lab. Chip 13, 4374–4384 (2013).

32. Nourmohammadzadeh, M. et al. A microfluidic array for real-time live-cell imaging of human and rodent pancreatic islets. Lab. Chip 16, 1466–1472 (2016).

33. Quintard, C., Achard, J.-L. & Fouillet, Y. Method for achieving microfluidic perfusion of a spheroid and device suitable for implementing said method. (2021).

34. Ajaev, V. S. & Homsy, G. M. Modeling shapes and dynamics of confined bubbles. Annu Rev Fluid Mech 38, 277–307 (2006).

35. Hathcock James J. Flow Effects on Coagulation and Thrombosis. Arterioscler. Thromb. Vasc. Biol. 26, 1729–1737 (2006).

36. Wimmer, R. A., Leopoldi, A., Aichinger, M., Kerjaschki, D. & Penninger, J. M. Generation of blood vessel organoids from human pluripotent stem cells. Nat. Protoc. 14, 3082–3100 (2019).

37. Zhang, S., Wan, Z. & D. Kamm, R. Vascularized organoids on a chip: strategies for engineering organoids with functional vasculature. Lab. Chip 21, 473–488 (2021).

38. Sudo, R. et al. Transport-mediated angiogenesis in 3D epithelial coculture. FASEB J. 23, 2155–2164 (2009).

39. Song, J. W., Bazou, D. & Munn, L. L. Anastomosis of endothelial sprouts forms new vessels in a tissue analogue of angiogenesis. Integr. Biol. 4, 857–862 (2012).

40. Nguyen, D.-H. T. et al. Biomimetic model to reconstitute angiogenic sprouting morphogenesis in vitro. Proc. Natl. Acad. Sci. 110, 6712–6717 (2013).

41. Oh, S. et al. “Open-top” microfluidic device for in vitro three-dimensional capillary beds. Lab. Chip 17, 3405–3414 (2017).

42. Vickerman, V. & Kamm, R. D. Mechanism of a flow-gated angiogenesis switch: early signaling events at cell–matrix and cell–cell junctions. Integr. Biol. 4, 863–874 (2012).

43. Carpentier, G., Martinelli, M., Courty, J. & Gascone, I. Angiogenesis analyzer for ImageJ. 4th ImageJ User and Developer Conference proceedings. (pp. 198–201). https://imagej.nih.gov/ij/macros/toolsets/Angiogenesis%20Analyzer.txt (2012).

## References

1. Bruus, H. Theoretical microfluidics. (Oxford University Press, 2007).

2. Shah, R. K. & London, A. L. Laminar Flow Forced Convection in Ducts. (1978).

3. Zivkovic, V., Zerna, P., Alwahabi, Z. T. & Biggs, M. J. A pressure drop correlation for low Reynolds number Newtonian flows through a rectangular orifice in a similarly shaped micro-channel. Chem. Eng. Res. Des. (2013) doi:10.1016/j.cherd.2012.05.022.

